# Herkogamy but not reciprocity is a better predictor of legitimate pollen transfer and fruit set in *Jasminum malabaricum*, a self-compatible species with stigma-height dimorphism

**DOI:** 10.1101/2020.06.11.145516

**Authors:** Shatarupa Ganguly, Deepak Barua

## Abstract

**Premise:** Reciprocity and herkogamy, morphological traits that define style length polymorphisms, are thought to be critical in determining legitimate inter-morph pollen transfer in plants with style length polymorphism. However, the consequences of individual-level variation in these traits for pollen transfer and reproductive success have rarely been examined, and the relationship between these two fundamental traits remains unexplored.

**Methods:** We quantified individual-level estimates of herkogamy and reciprocity and tested the assumption that higher herkogamy and reciprocity result in higher legitimate pollen transfer and reproductive success in natural populations of *Jasminum malabaricum*, a species that exhibits stigma-height dimorphism. Additionally, we examined the relationship between herkogamy and reciprocity to understand potential consequences for avoiding self-pollination and encouraging legitimate pollen deposition.

**Results:** Surprisingly, reciprocity was not related to pollen load, legitimate pollen fraction or reproductive success. In contrast, herkogamy was positively associated with legitimate pollen fraction and fruit set in the long-styled morph. Interestingly, we observed a negative relationship between herkogamy and reciprocity in the long-styled morph.

**Conclusions:** Herkogamy was more important than reciprocity in increasing legitimate pollen transfer and reproductive success in *J. malabaricum*. Herkogamy might be particularly important in stabilising species with stigma-height dimorphism and other such polymorphic intermediates with low reciprocity, and this may allow the evolution of reciprocal arrangement of sex organs at a later stage in the pathway towards distyly. The negative relationship between herkogamy and reciprocity suggests a trade-off between avoidance of self-pollen deposition and promotion of legitimate pollen deposition.

## Introduction

Reciprocity and herkogamy are the two defining morphological features in flowers of species with style length polymorphisms. Reciprocity is the physical match between complementary sex organ heights (Armbruster et al., 2017), while herkogamy is the separation of the male and female sex organs within a flower (Webb and Lloyd, 1986; Opedal, 2018). However, while reciprocity and herkogamy are widely assumed to play a critical role in the promotion of legitimate pollen transfer that is required to maintain style length polymorphisms, few studies have examined how variation in these traits is related to inter-morph pollen transfer and reproductive success in plants with style length polymorphisms (Cesaro et al., 2004; Thompson et al., 2012; Keller et al., 2014; Liu et al., 2016; Jacquemyn et al., 2018; Brys and Jacquemyn, 2020).

The reciprocal arrangement of sex organs in species with style length polymorphism enables pollinators to pick up pollen from anthers on specific locations on their body from where it can be accurately deposited on stigmas of the complementary morphs (Lloyd and Webb, 1992). Empirical studies in heterostylous species have focused on population-level estimates of reciprocity, and these have shown that higher reciprocity is related to greater legitimate pollen transfer (Lau and Bosque, 2003; Baena-Díaz et al., 2012), and reproductive success (Valois-Cuesta et al., 2011; Brys and Jacquemyn, 2014; Keller et al., 2014; Zhou et al., 2015; Ferrero et al., 2017). While limited, studies in species with stigma-height dimorphism have also shown that population-level estimates of reciprocity are positively related to reproductive success (Thompson et al., 2003, 2012; Stehlik et al., 2006). However, there are no studies that have examined individual level reciprocity in species with stigma-height dimorphism, and only two in species with heterostyly (Jacquemyn et al., 2018; Brys and Jacquemyn, 2020) that show that individuals with higher reciprocity have higher proportions of legitimate pollen.

The presence of self and heteromorphic incompatibility in most species with style length polymorphism makes it difficult to tease apart the contribution of reciprocity alone to legitimate pollination and maintenance of the polymorphism. Additionally, even in the presence of physiological incompatibility, reduced male and female fitness due to pollen discounting and stigma clogging, and associated reductions in reproductive success, have been observed (Shore and Barrett, 1984; Barrett and Glover, 1985; Martén-Rodríguez et al., 2013; Keller et al., 2014). In self-compatible species, higher legitimate pollen fractions do not guarantee a greater number of legitimate offspring. Substantial numbers of illegitimate offspring can result despite a higher legitimate fraction of pollen (Zhou et al., 2015). In such cases, the role of herkogamy in ensuring more legitimate offspring becomes crucial.

The function of herkogamy is to increase the fraction of legitimate pollen deposition on the stigma by reducing interference between male and female organs and reducing self and intra-morph pollen deposition (Webb and Lloyd, 1986; Barrett, 2002). Herkogamy has been studied extensively in monomorphic species that do not exhibit style length polymorphisms, and results from these have shown that greater stigma-anther separation between sex organs within flowers results in higher outcrossing rates (Ennos, 1981; Brunet and Eckert, 1998; Takebayashi et al., 2006; Larrinaga et al., 2009; de Vos et al., 2012, 2018; Li et al., 2013). However, empirical studies in species with style length polymorphisms are few, and these show that herkogamy is related to decreased self-pollination (Nishihiro and Washitani, 1998), as well as increased legitimate pollen transfer and reproductive success in heterostylous species (Liu et al. 2016). As with reciprocity, these studies focused on population-level estimates of herkogamy while ignoring intra-population variation in herkogamy (but see Hernandez and Ornelas 2007, Keller et al. 2014). Population-level estimates of herkogamy were also observed to be associated with decreased self-pollination in species with stigma-height dimorphism (Cesaro et al., 2004). Again, as with reciprocity, studies on consequences of intra-population variation in herkogamy lack in species with stigma-height dimorphism.

It is widely acknowledged that there can be a large variation in sex organ positions among individuals in populations with style length polymorphism, and this variation can have important consequences for reciprocity, herkogamy, pollen flow, and reproductive success for these individuals (Eckert and Barrett, 1994; Nishihiro et al., 2000; Sanchez et al., 2008; Sampson and Krebs, 2012; Costa et al., 2017). Population-level estimates of reciprocity and herkogamy can hide biologically important variation among individuals, and understanding intra-population variation in these traits is important as selection ultimately acts at the level of the individuals. Additionally, changes in organ positions in an individual flower have obvious implications for stigma-anther separation but are likely to also alter the match of these organs with counterparts in the complementary morph and affect reciprocity for that individual. The lack of an individual-level measure of reciprocity (Jacquemyn et al., 2018; Brys and Jacquemyn, 2020) and the focus on population-level estimates for both reciprocity and herkogamy have precluded such studies. Thus, the relationship between these two fundamental traits remain unexplored, and the consequences for the two distinct functions in avoiding self-pollen deposition and encouraging legitimate pollen deposition remain unknown.

In this study, we examined the variation in reciprocity and herkogamy among individuals of four naturally occurring isoplethic populations of *J. malabaricum* (Ganguly and Barua, 2020) with stigma-height dimorphism. We estimated reciprocity for an individual as the average mismatch between the stigma of that individual and the anthers of all other individuals of the complementary morph. We tested the assumption that an increase in reciprocity and herkogamy leads to a rise in legitimate pollen transfer and reproductive success, by examining the relationship of individual-level reciprocity and herkogamy with stigmatic pollen load and fruit set. Morph specific differences in stigmatic pollen load and reproductive success were also quantified to aid in the interpretation of the results as they can influence the above-mentioned relationships. We expected reciprocity to play an important role in legitimate pollen transfer and fruit set. However, given that our study species exhibits stigma-height dimorphism with low reciprocity and self-compatibility, we also expected herkogamy to significantly influence legitimate pollen transfer. Additionally, we examined the relationship between reciprocity and herkogamy across individuals to understand potential consequences for avoiding self-pollen deposition and encouraging legitimate pollen deposition.

## Methods

### Study sites and species

*Jasminum malabaricum* Wight. is a woody liana endemic to the Western Ghats of India, and Sri Lanka (Singh & Karthikeyan 2000) that exhibits stigma-height dimorphism (Ganguly, 2019). This study was conducted between 2015 and 2017 on four populations of *J. malabaricum:* namely, Trimbak (19.94° N, 73.54° E), Bhimashankar (19.07° N, 73.55° E), Mulshi (18.50° N, 73.51° E) and Kaas (17.72° N, 73.81° E). Spatiotemporal mating opportunities are similar for both morphs (Ganguly and Barua, 2020), and therefore, unlikely to be a confounding factor in our investigations.

### Quantification of herkogamy and reciprocity

One flower from 30 individuals of each morph was collected from Bhimashankar, Mulshi and Kaas, between March and April of 2015 and 2017, and preserved in formalin acetic alcohol (FAA; 2.5%:2.5%:95%) (Ganguly and Barua, 2020). In Trimbak, one flower was collected from 20 long-, and 19 short-styled individuals because of the small population size and limited availability of individuals. Herkogamy and reciprocity were also quantified for an additional 70 individuals for a total of 100 individuals of each morph in the Bhimashankar population. Anther and stigma heights were quantified using ImageJ (ver. 1.52a) from scanned images of dissected flowers. Since anthers were attached to the corolla tube, the distance from the base of the corolla tube to the mid-point of the anthers was measured as anther height. The distance from the base of the ovary to the mid-point of the stigma was measured as stigma height. Herkogamy was quantified for one flower per individual as the difference in anther and stigma height of a flower (Opedal, 2018). As we examined the relationship of reciprocity with measures of female fitness like pollen load and fruit set, reciprocity was calculated for one flower per individual as the mean of mismatches between stigma height for that individual and anther height of every other individual of the complementary morph. These pair-wise mismatches were averaged to give an estimate of reciprocity for that individual. We calculate mismatch, which is the inverse of reciprocity and hence higher values denote lower reciprocity. The relationship between herkogamy and reciprocity was examined using Pearson’s correlation coefficient. We use the term “mismatch” to refer to the spatial correspondence between anther and stigma heights of reciprocal morphs to describe our results. However, we use the term “reciprocity” to discuss the results to increase ease of interpretation.

### Quantification of stigma pollen load

We quantified stigma pollen load for the same individuals from the Bhimashankar (March 2016) and Kaas (April 2017) populations for which floral morphometry was quantified. Flowers of *J. malabaricum* open between 7 p.m. to 9 p.m. and last for approximately 33 h (Ganguly and Barua, 2020). Post 3 p.m. on the day after anthesis, the flowers wilt and no pollinator activity is seen on these flowers (Ganguly and Barua, 2020). Hence, to allow for maximal pollen deposition, flowers were collected between 2 and 3 p.m. the day after anthesis and preserved in formalin acetic alcohol (FAA; 2.5%:2.5%:95%). The quantification was done for 29 individuals of each morph for Bhimashankar, and 32 long-, and 31 short-styled individuals for Kaas. Stigmas were washed in 70% ethanol and softened in 4N NaOH for 12 h (Kearns and Inouye 1993). Subsequently, they were washed in distilled water and passed through serial concentrations of ethanol from 10% to 100% in steps of 15% to wash the oil from the pollen surface. The stigmas were then resuspended in a series of ethanol dilutions of glycerol with increasing concentration of glycerol from 10% to 90% in steps of 20%, squashed with a coverslip, and washed and mounted in 100% glycerol.

To quantify the total number of pollen deposited on a stigma, images of stigma squashes for two flowers per individual were taken at 40x magnification. An exhaustive pollen count was performed for all the stigmas. Legitimate pollen fraction was estimated for one stigma for each individual. Ten images were taken at 400x magnification for each stigma, and this accounted for greater than 15% of the total pollen on that stigma. Both the pollen count and measurement of pollen size were performed using ImageJ. As the equatorial and polar diameters were strongly correlated, only the equatorial diameter of the pollen was used as a measure of pollen size. The pollen were discriminated based on differences in diameter between the morphs using the estimates of pollen size obtained previously(Ganguly and Barua, 2020). The short-styled morph produced larger pollen, but pollen production was not significantly different between morphs. The diameter of the long-styled pollen ranged between 34 μm to 55 μm and the short-styled pollen between 47 μm to 73 μm (Ganguly and Barua, 2020). Approximately 35% of the pollen were in the range from 47 μm to 55 μm, which was common to both morphs. The ratio of the frequency of pollen grains extracted from anthers of the long- and short-styled morph in each size class of 1 μm was used to estimate the probability of occurrence of pollen from each morph in this overlapping size range. The pollen from the stigma which belonged to these size classes were distributed among the morphs based on this probability of occurrence. The legitimate pollen fraction was calculated as the percentage of the number of legitimate pollen identified in the total number of pollen sampled. Differences between morphs in natural total stigma pollen load and the legitimate pollen fraction were examined using a non-parametric Mann-Whitney U test as the values were not normally distributed.

### Autogamous pollen deposition

To investigate the function of herkogamy in decreasing self-pollination, three mature buds were tagged and emasculated for five individuals of each morph to quantify autogamous self-pollination in March 2017 in individuals from the Bhimashankar population. Another three buds on the same individuals were tagged as controls to account for individual differences in pollen deposition. Flowers were collected between 2 p.m. and 3 p.m. on the day after anthesis. The total pollen load and the legitimate fraction of pollen were quantified as described above. Wilcoxon’s signed-rank test was used to examine differences in total pollen load and the legitimate fraction between the control and the emasculated flowers from the same individual. This non-parametric test was chosen as the variables contained extreme values and so that control and treatment comparisons within the same individual could be made.

### Quantification of fruit set

Given the time-, and labour-intensive nature of this work, fruit set was quantified in March 2015 and 2016 in only one of the four study populations (Bhimashankar). This was done for the same 29 individuals of each morph for which we had quantified herkogamy and reciprocity and estimated stigma pollen load. For both morphs, 20 mature buds were tagged per individual in the peak flowering season and followed till the fruits matured. The percentage of total mature fruits formed from the 20 buds was taken as the final fruit set. Differences in natural fruit set between the two morphs were examined using a non-parametric Mann-Whitney U test as the values obtained were not normally distributed. The relationships of herkogamy and reciprocity with the natural total pollen load, the legitimate fraction of stigmatic pollen load and the fruit set were examined using Spearman’s rank correlation coefficient as most of the examined variables were counts and percentages, and had extreme values and skewed distributions.

## Results

There was substantial intra-population variation in estimates of mismatch and herkogamy between individuals in the populations (Fig. 1, Table 1). The long-styled morph exhibited a much larger range and higher coefficient of variation in mismatch as compared to the short-styled morph in all four study populations (Table 1). Similarly, the long-styled morph exhibited a larger range and higher coefficient of variation in herkogamy as compared to the short-styled morph (Table 1).

**Figure 1:**
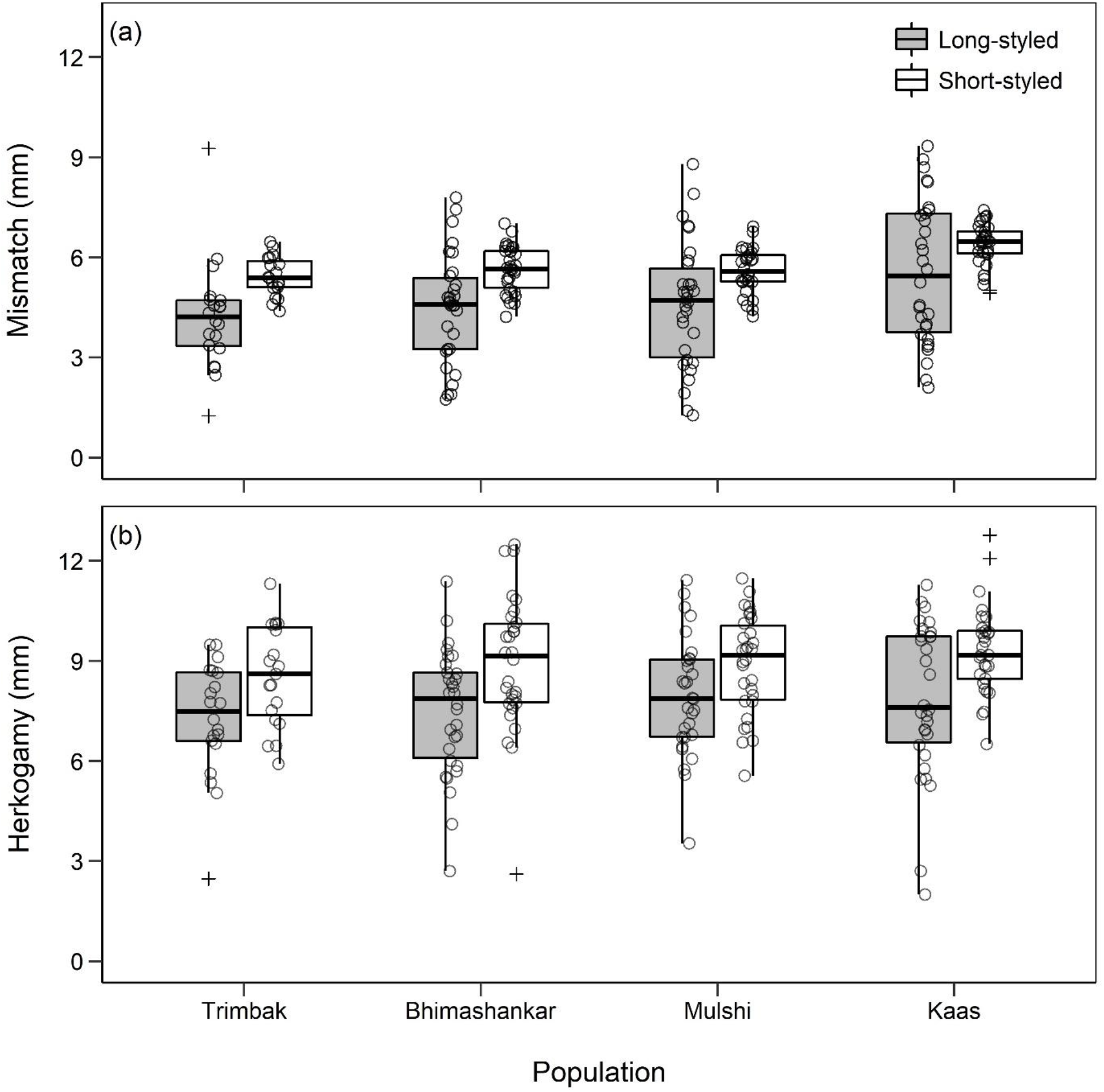
(a) Mismatch and (b) herkogamy in the long-styled (dark grey boxes) and short-styled (open boxes) morphs of the four study populations. The number of individuals sampled for the long- and the short-styled morph for the populations are 20 and 19 for Trimbak; 30 and 30 for Bhimashankar, Mulshi, and Kaas. Outliers are denoted by +. The bold line in the boxplots represents the median, the ends of the box represent the first and the third quartiles and the whiskers represent 1.5 times the interquartile range.

**Table 1:** Range (Min-Max) and coefficient of variation (CV) of individual mismatch and herkogamy observed in the four study populations a) Trimbak, b) Bhimashankar, c) Mulshi and d) Kaas. L and S stand for long- and short-styled morph, respectively. The number of individuals sampled for the long- and the short-styled morph are 20 and 19 for Trimbak; 30 and 30 for Bhimashankar, Mulshi, and Kaas.

| Population | Morph | Mismatch |  | Herkogamy |  |
| --- | --- | --- | --- | --- | --- |
|  |  | Min-Max (mm) | CV % | Min-Max (mm) | CV % |
| a) Trimbak | L | 1.25, 9.27 | 38.70 | 2.47, 9.49 | 23.90 |
|  | S | 4.39, 6.47 | 10.80 | 5.91, 11.30 | 17.70 |
| b) Bhimashankar | L | 1.74, 7.80 | 36.70 | 2.70, 11.40 | 25.20 |
|  | S | 4.22, 7.02 | 12.30 | 2.61, 12.50 | 22.90 |
| c) Mulshi | L | 1.27, 8.80 | 41.10 | 3.53, 11.40 | 22.20 |
|  | S | 4.24, 6.92 | 12.00 | 5.56, 11.50 | 16.80 |
| d) Kaas | L | 2.10, 9.34 | 38.50 | 2.00, 11.30 | 29.60 |
|  | S | 4.93, 7.42 | 10.50 | 6.51, 13.10 | 15.80 |

There was high variation between individuals in natural total stigma pollen load, with counts ranging from 13 to 5015 in the long-styled, and 38 to 1394 in the short-styled morph in the Bhimashankar population, and from 6 to 1542 in the long-styled and 1 to 1350 in the short-styled morph (Fig. 2 a, c) in the Kaas population. Although long-styled morphs had higher mean stigma pollen load, no significant differences were detected between the two morphs in either of the two populations examined (Fig. 2 a, c; Mann-Whitney U test: Bhimashankar *p* = 0.41 and Kaas *p* = 0.11; *n* = 29 and 29 individuals for Bhimashankar, and 32 and 31 individuals for Kaas for the long- and short-styled morphs). The legitimate pollen fraction was significantly higher in the short-than in the long-styled morph for both populations (Fig 2 b, d; Mann-Whitney U test: Bhimashankar *p* < 0.001 and Kaas *p* < 0.001; *n* for long-styled and short-styled morph was 29 and 29 individuals for Bhimashankar, and 32 and 31 individuals for Kaas).

**Figure 2:**
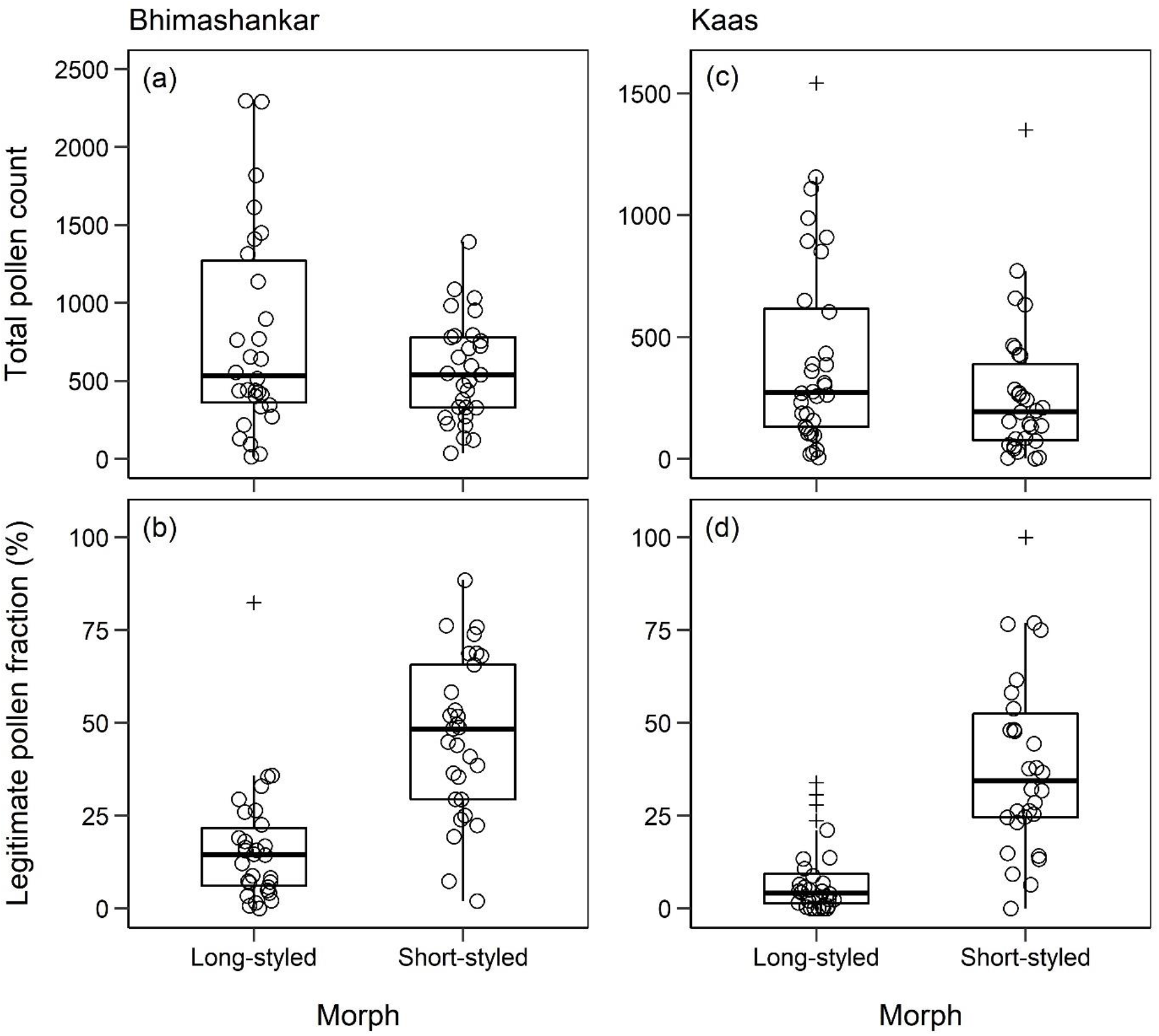
The total stigma pollen load (a, c) and the legitimate pollen fraction (b, d) in the Bhimashankar and Kaas populations. The number of individuals sampled for the long- and short-styled morph is 29 and 29 for Bhimashankar, and 32 and 31 for Kaas. One extreme outlier value of greater than 5000 for total pollen load in a long-styled individual from the Bhimashankar population has been removed from the plot for better visualisation of the differences between the morphs. The bold line in the boxplots represents the median, the ends of the box represent the first and the third quartiles and the whiskers represent 1.5 times the interquartile range.

In both morphs, emasculation resulted in a decrease in the total pollen deposition (Fig. 3 a; Wilcoxon signed-rank test: long-styled *p* = 0.04 and short-styled *p* = 0.04; 3 flowers per treatment for 5 individuals of each morph). In the short-styled morphs emasculation did not affect the legitimate pollen fraction (Fig. 3 b; Wilcoxon signed-rank test: *p* = 0.34). In contrast, in the long-styled morphs emasculation resulted in an increase in the legitimate pollen fraction (Wilcoxon signed-rank test: *p* = 0.04). Fruit set was not significantly different between the long-, and short-styled morphs and this was consistent across the two years in which we examined this in individuals from the Bhimashankar population (Fig. 4; Mann-Whitney U test: 2015 *p* = 0.41; 2016 *p* = 0.80; *n* = 29 individuals each morph in both years).

**Figure 3:**
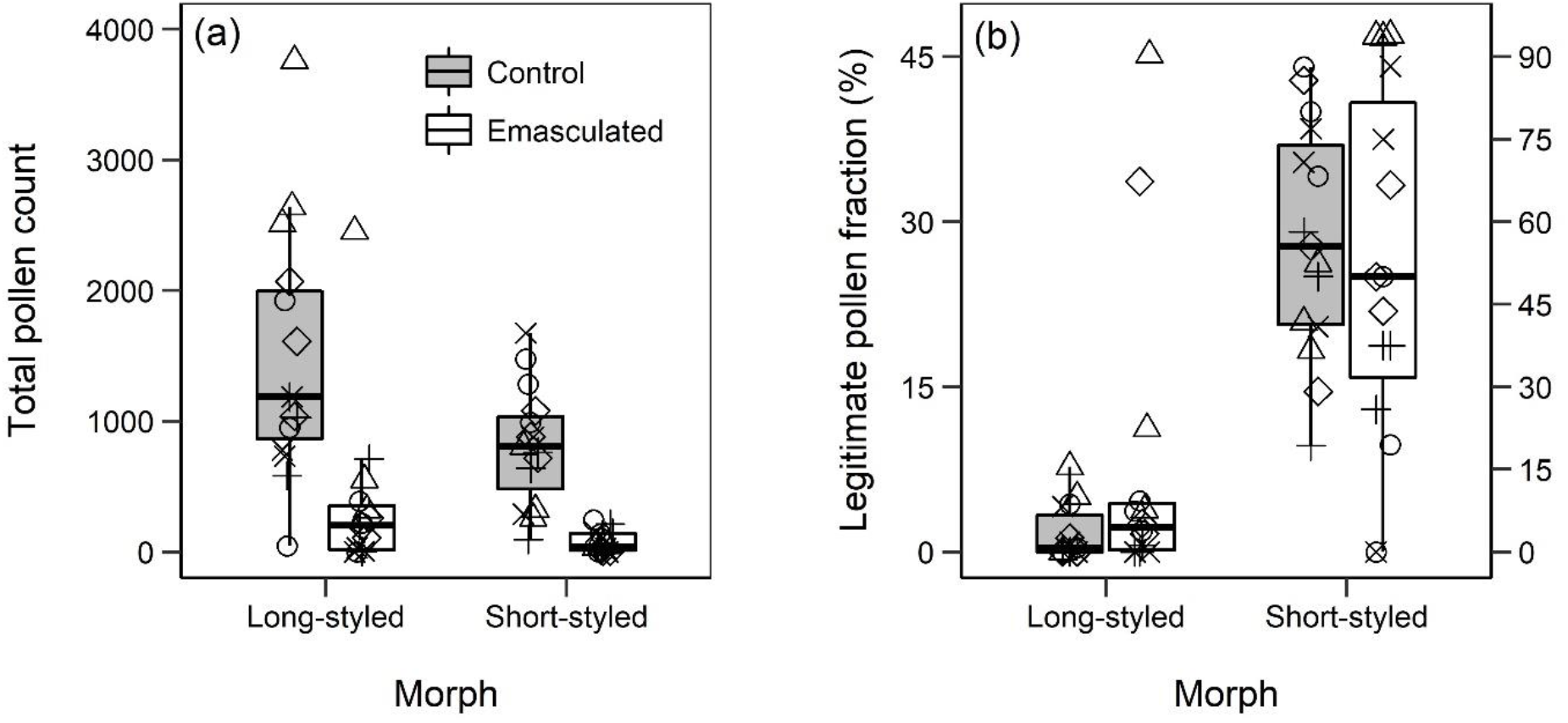
(a) The total pollen load, and (b) the legitimate pollen fraction in control (non-emasculated, grey boxes) and emasculated flowers (open boxes) to determine autogamous pollen deposition. One point from the legitimate pollen fraction for the emasculated group of long-styled morph (80%) has been removed from this plot for better visualisation of the differences between the control and emasculation treatments. Data are from three flowers from five individuals (represented by the different symbols). The bold line in the boxplots represents the median, the ends of the box represent the first and the third quartiles and the whiskers represent 1.5 times the interquartile range.

**Figure 4:**
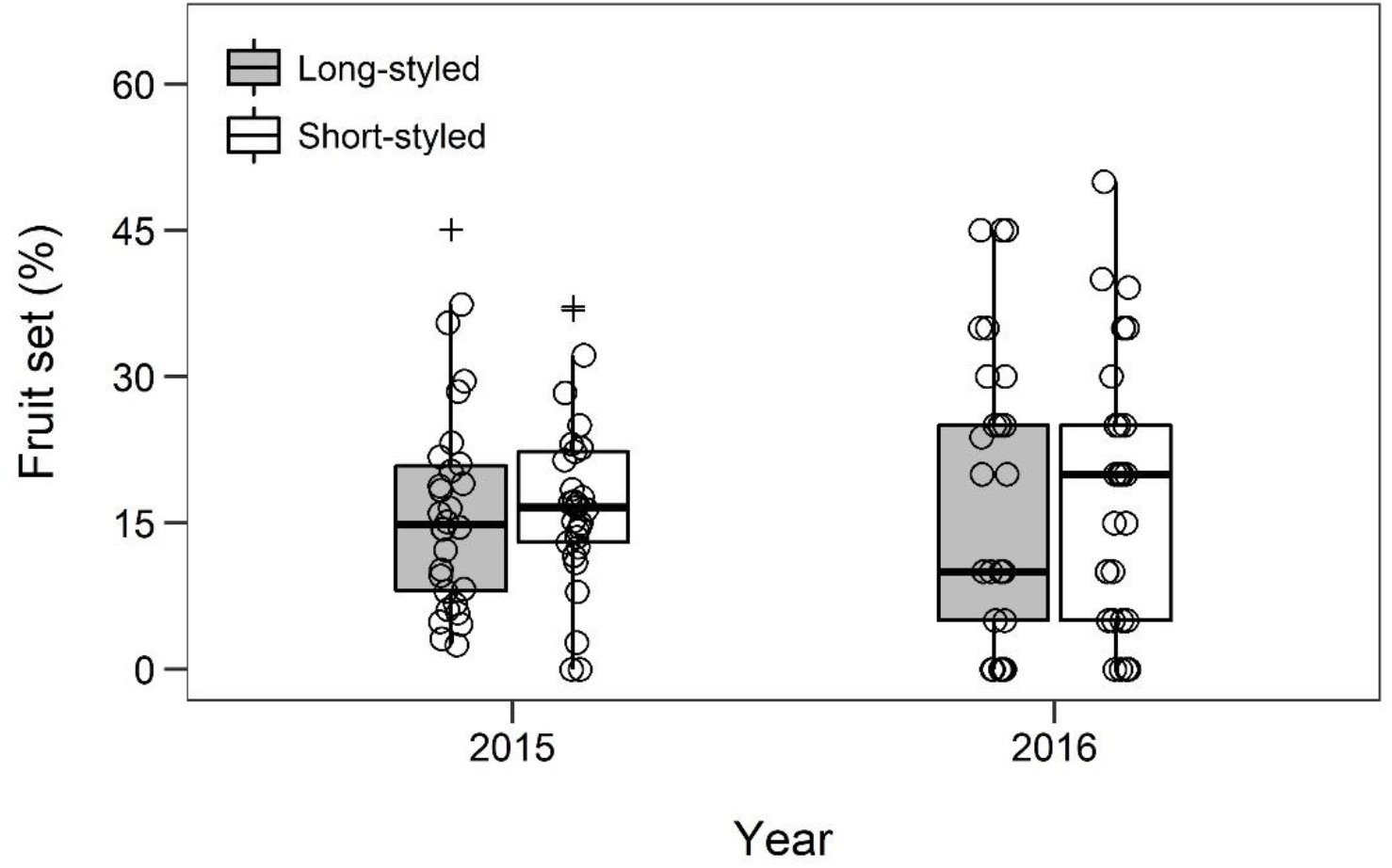
Natural fruit set in long- (grey boxes) and short-styled (open boxes) individuals from the Bhimashankar population measured as the percentage fruit formed per flower in 2015 and 2016. The number of individuals sampled for the long- and the short-styled morph are 29 and 29 respectively. The plus sign represents outliers. The bold line in the boxplots represents the median, the ends of the box represent the first and the third quartiles and the whiskers represent 1.5 times the interquartile range.

Mismatch was not related to total pollen deposition or the legitimate pollen fraction. Counter-intuitively, mismatch was positively related to fruit set in the short-styled morph in Bhimashankar (Table 2 a). Herkogamy, on the other hand, was positively related to disassortative pollen load and fruit set in the long-styled morph, but not in the short-styled morph in Bhimashankar (Table 2 a). Herkogamy was negatively related to total pollen load in the long-styled morph, but not related to pollen deposition in the short-styled morph in Kaas (Table 2 b).

**Table 2:** Relationship of herkogamy and reciprocity to total stigma pollen load, legitimate pollen fraction and fruit set in the Long-styled (L) and short-styled (S) morph in a) Bhimashankar, and b) Kaas. Fruit set was estimated only for the Bhimashankar population. *P* < 0.05 and *P* < 0.1 are denoted by * and ^ψ^, respectively, next to the Spearman’s rank-order correlation coefficients presented. The number of individuals sampled for the long- and short-styled morph for the two populations is 29 and 29 for Bhimashankar, and 32 and 31for Kaas.

| Population | Trait | Morph | Total pollen load | Legitimate fraction | Fruit set |
| --- | --- | --- | --- | --- | --- |
| a) Bhimashankar | Mismatch | L | 0.08 | 0.23 | 0.28 |
| | | S | 0.03 | -0.22 | 0.33Ψ |
|  | Herkogamy | L | 0.11 | 0.43* | 0.42* |
|  |  | S | -0.11 | -0.17 | 0.01 |
| b) Kaas | Mismatch | L | -0.18 | 0.04 |  |
|  |  | S | 0.19 | -0.18 |  |
| | Herkogamy | L | -0.34Ψ | 0.14 | |
|  |  | S | -0.27 | -0.04 |  |

Herkogamy and mismatch showed a significant positive correlation in the long-styled morph in all study populations (Fig. 5 a-d). However, such a positive relationship between herkogamy and mismatch was seen in only one of the four study populations in the short-styled morph (Fig. 5 e-h). Similar results were observed when examining this relationship in a larger number of individuals from the Bhimashankar population with a significant positive relationship between herkogamy and mismatch in the long-, but not the short-styled morph (Supplementary Information Fig. A1 a, b).

**Figure 5:**
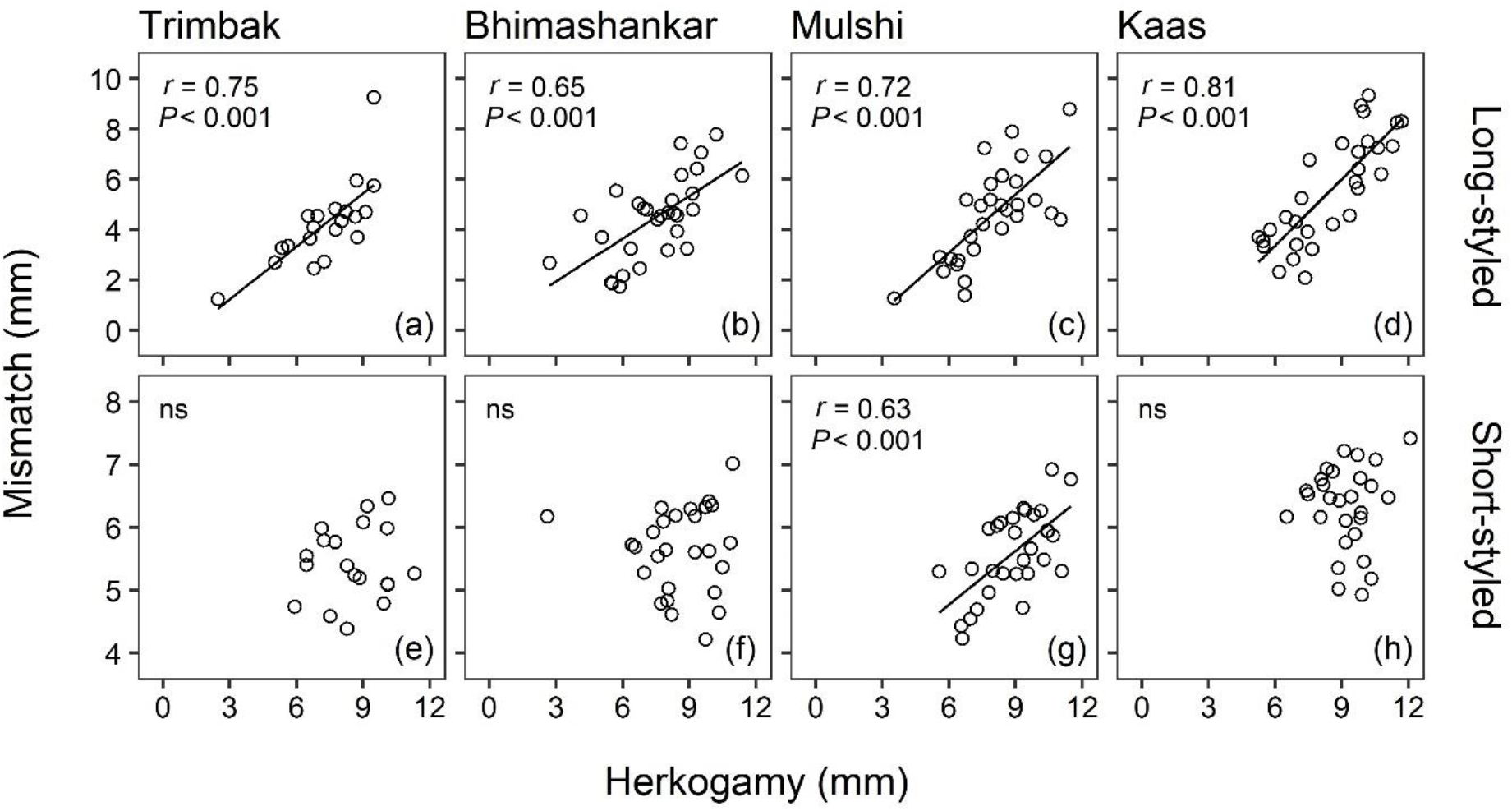
Relationship between herkogamy and mismatch in the two morphs (long-styled morph: a,b,c,d; short-styled morph: e,f,g,h) in the four study populations. Pearson’s correlation coefficient (*r*), and the corresponding *P-*value are shown where *P* < 0.05, and ‘ns’ denotes not significant. The number of individuals sampled for the long- and the short-styled morph for the populations are 20 and 19 for Trimbak; 30 and 30 for Bhimashankar, Mulshi, and Kaas.

## Discussion

Surprisingly, individual-level variation in reciprocity was not related to legitimate pollen fraction or fruit set. In contrast, individuals of the long-styled morph with higher herkogamy had a higher fraction of legitimate pollen and greater fruit set. Thus, herkogamy may be relatively more important than reciprocity in increasing legitimate pollen deposition and reproductive success in *Jasminum malabaricum*, a species with stigma-height dimorphism that lacks self-, and heteromorphic incompatibility. Additionally, we report here for the first time a negative relationship between reciprocity and herkogamy, the two defining traits of style length polymorphism. This suggests a potential trade-off between increasing legitimate pollen deposition and avoidance of self-pollen deposition.

Intra-population variation in anther and stigma heights is expected to reduce reciprocity and hence legitimate pollen transfer and reproductive success in species with style length polymorphism (Sanchez et al., 2008; Armbruster et al., 2017). We observed large variation in anther and stigma positions, and in the resultant reciprocity in individuals of all populations of our study species. However, this observed variation in reciprocity was not associated with any negative consequences and was unrelated to the legitimate fraction of pollen load or fruit set. Previous studies have shown that higher herkogamy but not reciprocity was related to higher legitimate pollen transfer and reproductive success (Keller et al., 2014). Additionally, legitimate pollen transfer in some species with style length polymorphism is likely to be primarily governed by effective long-tongued pollinators rather than sex organ positions (Simón-Porcar et al., 2014).

The higher fraction of legitimate pollen received by the short-styled morph is likely due to reduced self-pollen deposition as a consequence of higher herkogamy in this morph (Liu et al., 2016). High legitimate pollen transfer has also been associated with efficient long-tongued pollinators (Haddadchi, 2013; Simón-Porcar et al., 2014) that are capable of reaching the short-styled stigma. However, the higher legitimate pollen fractions observed in the short-styled morph did not translate into higher fruit set. As both morphs lack physiological incompatibility, similar total pollen loads resulted in equal fruit set between morphs. This is congruent with an observation from a previous study, where despite much higher legitimate pollen deposition, a considerable number of illegitimate offspring are formed due to the absence of self- and heteromorphic incompatibility (Zhou et al., 2015).

In the long-styled morph, emasculation resulted in a decrease in total pollen load and an increase in the legitimate pollen fraction. Thus, reduced self pollen deposition with increasing herkogamy is likely to result in the negative relationship between herkogamy and total pollen load, and the positive relationship between herkogamy and the legitimate pollen fraction that was observed in the long-styled morph. The higher fruit set associated with higher herkogamy in the same morph could be attributed to lower mechanical interference between the male and female sex organs. The lack of any change in the legitimate pollen fraction with emasculation in the short-styled morph implies that the higher herkogamy observed in this morph was sufficient to avoid self-pollen deposition. This was also reported in *Persicaria japonica* where emasculation had no effect on the deposition of incompatible pollen (Nishihiro and Washitani, 1998). This likely explains the absence of a relationship between herkogamy and reproductive success in the short-styled morph, where due to the higher herkogamy observed in this morph minor changes in herkogamy did not influence self-pollen deposition or fruit set. Similar results were reported in *Primula* spp., where the legitimate pollen fraction increased with herkogamy in the long-, but not the short-styled morph, and the “threshold” effect as mentioned above was offered as a possible explanation (Keller et al., 2014).

Reciprocity was negatively related to herkogamy in individuals of the long-styled morph. This relationship suggests a potential trade-off between maximising legitimate pollen transfer and avoiding self-pollination and will be useful in providing insights into the response of these species under different ecological scenarios. For example, in situations of pollen limitation due to the scarcity of mate and pollinators, and an absence of intra-morph incompatibility or inbreeding depression (Baker et al., 2000), reproductive assurance is important. If the relationship between reciprocity and herkogamy is negative, the individuals with high reciprocity and low herkogamy will be selected for as they would have the highest reproductive assurance in the population. A positive relationship can be maladaptive in such a situation. Hence, such relationships can help us understand the change in the distribution of anther and stigma heights in a population under different ecological scenarios. There was no relationship between reciprocity and herkogamy in individuals of the short-styled morph with the exception of the Mulshi population where, similar to the long-styled individuals, a negative relationship was observed. The difference in the relationship between reciprocity and herkogamy between morphs and among populations indicates the effect of morph specific selection pressures and differences in local environmental factors among populations (Santos-Gally et al., 2013). The short-styled morph does not show a relationship even at higher sample size, implying that the lack of a relationship is unlikely due to insufficient sample sizes.

Several studies have highlighted the functional significance of sex organ positions in species with style length polymorphism (Faivre and McDade, 2001; Sosenski et al., 2010; Ferrero, Castro, et al., 2011; Ferrero, Chapela, et al., 2011; Keller et al., 2012; Yuan et al., 2017), but the emphasis of these has primarily been on reciprocity at the level of populations while herkogamy has been relatively understudied (but see Nishihiro and Washitani 1998, Keller et al. 2014, Liu et al. 2016, Faife-Cabrera et al. 2016, Barranco et al. 2019). Moreover, although it has been acknowledged that intra-population variation in sex organ positions affects legitimate pollen transfer and reproductive success (Eckert and Barrett, 1994; Lau and Bosque, 2003; Sanchez et al., 2008; Armbruster et al., 2017), studies examining this are rare. This study reveals that herkogamy can be more important than reciprocity in promoting legitimate pollen transfer and fruit set. The result highlights the functional role of herkogamy and has especially important implications for the maintenance of imperfectly reciprocal intermediates in the pathway towards heterostyly. The results also suggest that intra-population variation in herkogamy and reciprocity might not affect legitimate pollen transfer and reproductive success across individuals and is perhaps the reason why species with style length polymorphisms display high variation in sex organ positions. Examination of the intra-population variation in reciprocity and herkogamy also allowed us to investigate the relationship between the two, which revealed a trade-off between self-pollination and legitimate pollination. Further studies on the causes and consequences of intra-population variation in reciprocity and herkogamy can substantially increase our understanding of the coevolution of the dimensions of floral parts with pollinators.

## Author contributions

SG and DB conceived the idea. SG conducted field studies to collect the data. SG analysed the data. SG and DB wrote the manuscript.

## Acknowledgements

We thank the Maharashtra State Biodiversity Board and the Maharashtra Forest Department for permits to work on *J. malabaricum*; Kalu Kurade, Ganpat Lokhare and Thakku Kurade for assistance in the field; Neha Wadmare for help in quantifying floral morphology; Rashmi Rai and Zakhiya P.C. for help in pollen size quantification; and Yashraj Chavhan, Souparna Chakrabarty and Nitish for occasional assistance in fieldwork. This research was supported by intramural funding from the Indian Institute of Science Education and Research Pune (IISER Pune, Maharashtra, India).

