## Supplementary Figure 1 for "Herkogamy but not reciprocity is a better predictor of legitimate pollen transfer and fruit set in *Jasminum malabaricum*, a self-compatible species with stigma-height dimorphism"

**Supplementary material**

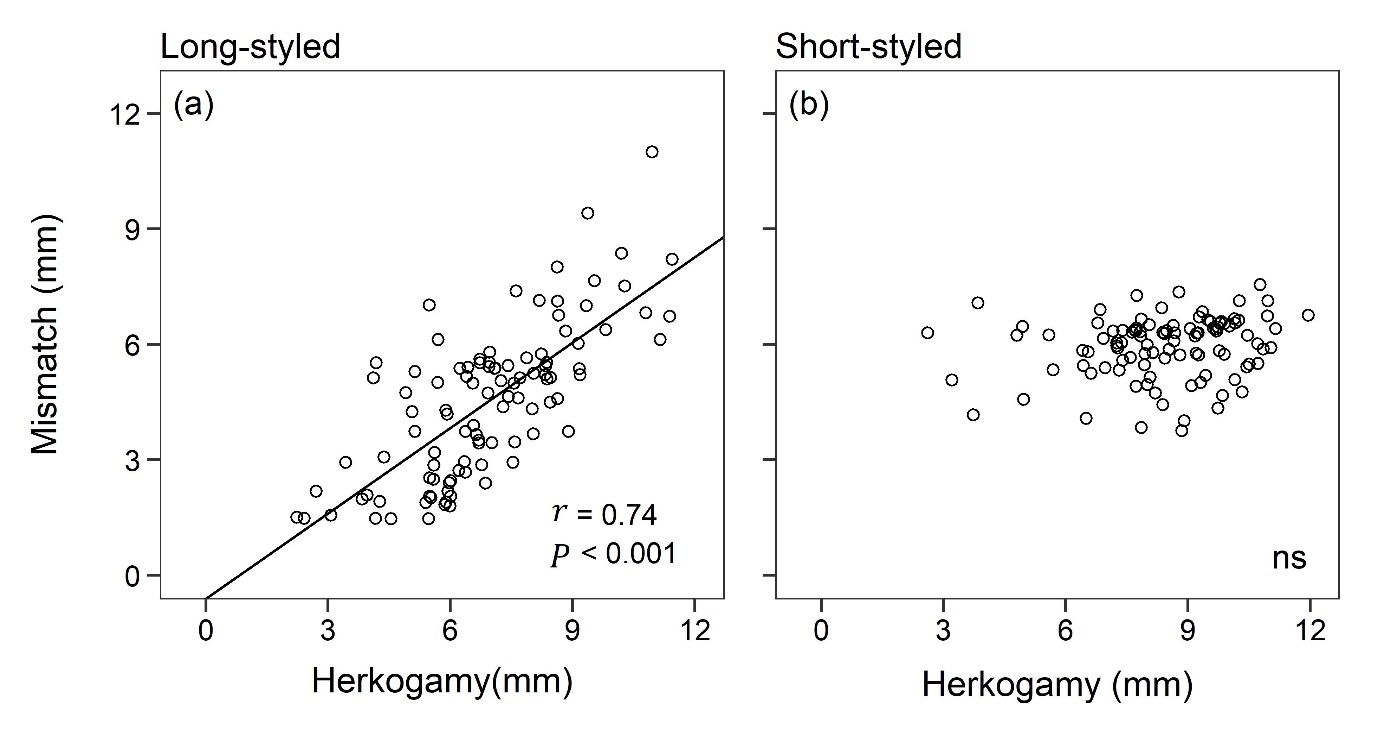

**Figure S1:** Relationship between herkogamy and mismatch in the (a) long-styled morph and (b) short-styled morph for individuals from Bhimashankar. Pearson's correlation coefficient (*r*) and the corresponding *P-*value are shown where *P* < 0.05, and ‘ns’ denotes not significant. A total of 100 individuals each were sampled for the long- and the short-styled morph.
